# Enhanced translation of leaderless mRNAs under oxidative stress in *Escherichia coli*

**DOI:** 10.1101/2021.06.29.449897

**Authors:** Lorenzo Eugenio Leiva, Omar Orellana, Michael Ibba, Assaf Katz

**Author notes:** **Correspondence**: Assaf Katz.

## Abstract

The bacterial response to oxidative stress requires the adaptation of the proteome to the hostile environment. It has been reported that oxidative stress induces a strong and global inhibition of both, transcription and translation. Nevertheless, whereas it is well known that transcription of a small group of genes is induced thanks to transcription factors such as OxyR and SoxR, an equivalent mechanism has not been described for translation. Here we report that whereas canonical translation that depends on Shine Dalgarno recognition is inhibited by oxidative stress in *Escherichia coli*, the translation of leaderless mRNA (lmRNA) is enhanced under such conditions. Both, inhibition of canonical translation and enhancement of lmRNA translation, depend on the production of (p)ppGpp. We propose that such a mechanism would allow bacteria to rapidly adapt their proteome to hostile conditions and is, perhaps, a general strategy to confront strong stressful conditions.

**Significance statement:** The regulation of translation (the production of proteins based on genetic information) is central for the adaptation to environmental changes. In *Escherichia coli* translation may begin through two alternative pathways. 1.- A canonical initiation that is well understood and is regulated mostly by changes in the accessibility of ribosomes to specific sequences and 2.- Initiation of leaderless mRNAs (lmRNAs) that lack these sequences and for which we do not understand the regulation process. Our results indicate that under oxidative stress, the production of (p)ppGpp in *E. coli* inhibits canonical translation and simultaneously enhances translation of lmRNAs, showing for the first time a natural condition where lmRNA translation is regulated and a role for (p)ppGpp in this process.

## Introduction

Bacteria are exposed to continual environmental changes, where modification of the proteome allows adaptation to the new conditions. Regulation of bacterial translation is key to this process (Rothschild & Mancinelli, 2001) and usually depends on changes to the initiation step. This is usually the slowest step of translation, and thus, changes to it will produce the greatest effects on protein production (Gualerzi & Pon, 2015; Laursen et al., 2005). According to the canonical model, initiation of translation in bacteria starts with the positioning of the small ribosomal subunit (30S) and initiator tRNA (fMet-tRNA) at the start codon in the mRNA, in conjunction with initiator factors 1 (IF1), 2 (IF2) and 3 (IF3), generating a pre-initiation complex (Gualerzi & Pon, 2015; Schmeing & Ramakrishnan, 2009). The region located upstream of the initiation codon, the leader region, guides the positioning of the start codon in the peptidyl-tRNA site (P-site) of the 30S subunit, by the interaction between the Shine Dalgarno (SD) sequence in the mRNA and a highly conserved segment of the 3’ end of 16S rRNA called anti-SD (aSD) sequence. In addition, low folding stability (Saito et al., 2020) and interactions of the ribosomal protein S1 and enhancer sequences such as a A/U-rich region upstream of SD may aid binding and positioning of the mRNA to the small ribosomal subunit (Kaminishi et al., 2007; Komarova et al., 2002). Positioning of the 30S at the ribosome binding site (RBS) allows binding of the large ribosomal subunit (50S), hydrolysis of GTP by IF2 and the release of initiation factors. This enables the entry of the elongation factor Tu (EF-Tu) with aminoacylated tRNAs and the start of polypeptide synthesis (Gualerzi & Pon, 2015; Schmeing & Ramakrishnan, 2009). Given the relevance of the positioning of the 30S subunit on the RBS, any change of its accessibility will alter the speed of translation initiation. Thus, changes of RNA folding or binding of sRNA and proteins have been observed to be usually involved in translation regulation (Duval et al., 2015).

In addition to the canonical initiation of translation, bacteria may initiate translation from leaderless mRNA (lmRNA). These mRNA are characterized by its lack of the usual 5’ UTR. They may present none or a small number of nucleotides and, most importantly, they completely lack any SD sequence. In contrast to the canonical initiation of bacterial translation, lmRNA translation has been proposed to start by binding of 70S or smaller 61S particles, not 30S (Beck et al., 2016; O’Donnell & Janssen, 2002; Udagawa et al., 2004). These kind of mRNA without a canonical RBS have been found in phages (e.g., λ and P2), transposons (e.g., Tn1721), and several bacterial species (e.g., *Mycobacterium smegmatis* and *Streptomyces clavuligerus*) (Cortes et al., 2013; de Groot et al., 2014; Hwang et al., 2019; T. G. Nguyen et al., 2020; Resch et al., 1995). Although they are common in nature, studies targeted to describe transcription initiation sites have usually found only a small number of lmRNA in enterobacteria. For instance, only 20 to 30 lmRNA were found in *E. coli* K-12 and *Salmonella enterica* serovar Typhimurium SL1344 (Kröger et al., 2012; Mendoza-Vargas et al., 2009). Nevertheless, it has been reported in other enterobacteria that transcription from alternative promoters may allow the production of lmRNA under hostile conditions. For example, an alternative promoter has been described for *virF*, a transcriptional factor that regulates virulence of *Shigella* spp. during mammalian cell infection. This alternative promoter generates a lmRNA that allows the production of a functional fragment of the *virF* regulator (Di Martino et al., 2016), suggesting a possible involvement of lmRNAs translation in the adaptation to host-pathogen conditions.

### Translation under oxidative stress

An example of the hostile conditions commonly confronted by bacteria is oxidative stress. Bacteria are often exposed to various environmental and anthropogenic factors that favor the production of oxidative molecules, which in some conditions, can produce oxidative stress (Imlay, 2019; Kataria & Ruhal, 2014). One of such conditions is the interaction with human immune cells, such as macrophages and polymorphonuclear leukocytes (G. T. Nguyen et al., 2017; Slauch, 2011). As a consequence, microorganisms have developed diverse strategies to overcome this stress. Whereas the transcriptional regulation of this response has been described in great detail (Chiang & Schellhorn, 2012; Imlay, 2013), the literature on changes in the translational machinery under oxidative stress is more limited. However, many authors have described a global inhibition of translation as a consequence of oxidative stress (Fasnacht & Polacek, 2021; Kojima et al., 2007; Nishiyama et al., 2004; Shenton et al., 2006; Zhong et al., 2015; M. Zhu & Dai, 2019). In addition to this global repression of translation, we and other authors have described important alterations in translation elongation. For instance, an increased error rate in the incorporation of some amino acids (Ling & Soll, 2010; Steiner et al., 2019; Wu et al., 2014) or decreased elongation rates during translation have been observed (Leiva et al., 2020; Zhong et al., 2015; M. Zhu & Dai, 2019). The relative contribution of each of these effects is not completely understood. Some groups have proposed that tRNA degradation limits translation elongation, determining the global translation efficiency (Zhong et al., 2015; M. Zhu & Dai, 2019). Nevertheless, this effect depends on the stressor concentration and is probably affected by other aspects, such as culture conditions and particular strains used. In effect, we have not observed this tRNA degradation using another strain of *E. coli* (K-12 MG1655). When using 250 μM of paraquat to induce oxidative stress, the decreased translation seems to derive mainly from steps different from elongation, probably translation initiation (Leiva et al., 2020).

To better understand the impact of oxidative stress on translation, here, we have investigated the impact of this stress on translation initiation. Our results show that while translation of canonical mRNA is inhibited under oxidative stress, that of lmRNAs is strongly induced. Furthermore, the results strongly suggest that these changes depend in part on the production of (p)ppGpp, an alarmone known to modulate the transcription and translation machinery in response to hostile environmental conditions.

## Results

### Oxidative stress induces the formation of a ribosomal particle of lower sedimentation coefficient

In order to get some insights about the effects of paraquat on translation, we analyzed the sedimentation coefficient of ribosomes from *E. coli* cultured in M9r media in the presence of 250 μM paraquat. This is a technique commonly used to analyze translation, as it indicates the proportion of ribosomes that are involved in more efficient (polysome fraction) and more inefficient (monosome/70S fraction) translation. Previous assays have shown that under the oxidative stress condition used in this experiment, bacterial replication and protein synthesis are greatly decreased (Leiva et al., 2020). Also, for this strain, the decreased translation produced by this lower concentration of paraquat seems to depend mainly on steps different from elongation (Leiva et al., 2020). To this end, a sedimentation analysis of the components of the translational machinery was carried by ultracentrifugation in sucrose gradients (10-40%) (Qin & Fredrick, 2013). As expected, free 30S, and 50S subunits, a majority peak corresponding to monosomes (70S) and a fraction of polysomes were observed when analyzing cells cultured under control conditions (Figure 1A) (Qin & Fredrick, 2013). These peaks were also observed in extracts from cultures stressed with 250 μM paraquat for 30 minutes (Figure 1B). However, we observed the appearance of a particle of intermediate sedimentation coefficient between the 50S subunit and complete ribosome (70S) peaks, whose abundance varies over time (Figure 1C). By interpolation, we have estimated that this particle has a sedimentation coefficient of 60 ± 0.6S, very similar to particles observed after addition of Kasuganycin to *E. coli* cultures, especially when lmRNAs are overexpressed. This antibiotic has been proposed to promote the loss of at least 6 ribosomal proteins of the minor subunit (some of them highly relevant for translation, such as S1 and S12) and induce the translation of lmRNA (Delvillani et al., 2011; Kaberdina et al., 2009). Given the relevance that activation of translation of the lmRNA subset of the transcriptome would have in the adaptation to stressful conditions, we decided to further investigate the possibility that translation of lmRNA is induced by paraquat dependent oxidative stress.

**Figure 1.**
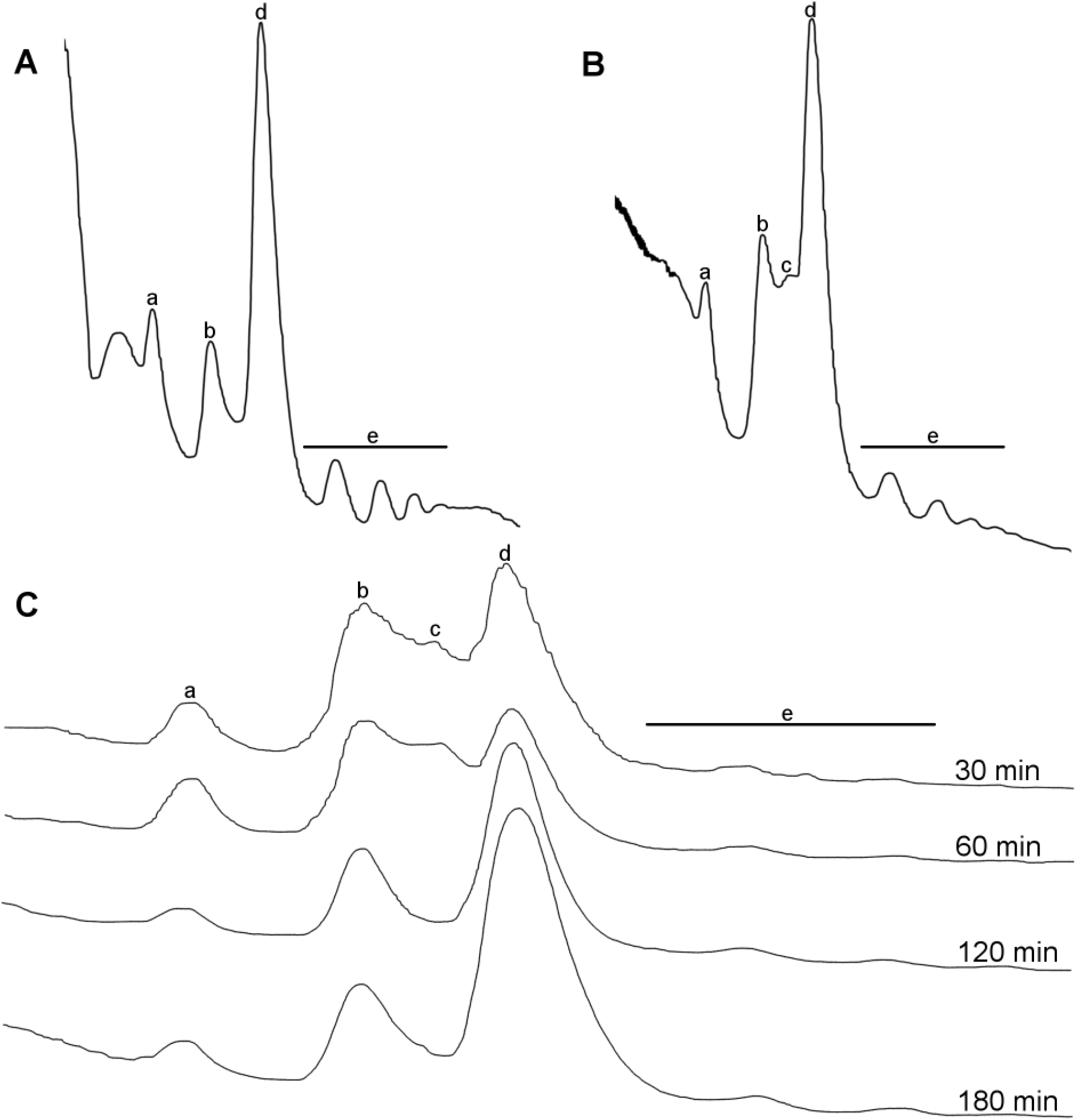
Oxidative stress induces the appearance of an undescribed peak between 50S and 70S. Sedimentation profiles of *E. coli* extracts separated in 10 to 40% sucrose gradients and detected at 254 nm. A) Samples collected from cultures at control conditions and B) 30 min oxidative stress (Paraquat 250μM). C) Similar experiments performed after 30, 60, 120 and 180 min of oxidative stress induced by 250 μM Paraquat. Lowercase letters indicate 30S (a), 50S (b), new peak (c), 70S (d) and polysomes (e).

### Translation of leaderless mRNA is enhanced by oxidative stress

Canonical and lmRNA initiation are the only two mechanisms known to be capable of initiating translation of the first cistron of bacterial transcripts. To compare the efficiency of translation by both mechanisms under control and oxidative stress conditions, we used two genetic reporter plasmids that code for a biscistronic operon coding for two fluorescent proteins, GFP upstream of mCherry (Figure 2A). One of such plasmids, plasmid pSD, was used to study canonical initiation. In pSD, both reporter genes (*gfp* and *mCherry*) have a canonical RBS. A second plasmid, plmRNA, was used to study lmRNA initiation. In plmRNA the first cistron (*gfp*) lacks a leader region, but the second cistron (mCherry) preserves a canonical RBS. Transcription initiation sites for both reporter plasmids were confirmed by reverse transcription coupled to template exchange (Langevin et al., 2013; Y. Y. Zhu et al., 2001). These experiments indicated that the lmRNA transcript presents 3 nucleotides (ATA) upstream of the AUG initiation codon confirming its leaderless nature (Supplementary Table S1). Thus, in both plasmids mCherry initiation operates through the same canonical mechanism, but while initiation of *gfp* translation depends on canonical mechanisms for pSD, in plmRNA *gfp* translation may only start through leaderless translation. As both genes, *gfp* and *mCherry*, are coded in the same RNA, alterations in the ratio of fluorescence measured from both proteins (GFP/mCherry) will reflect relative changes in the efficiency of both translation mechanisms.

**Figure 2.**
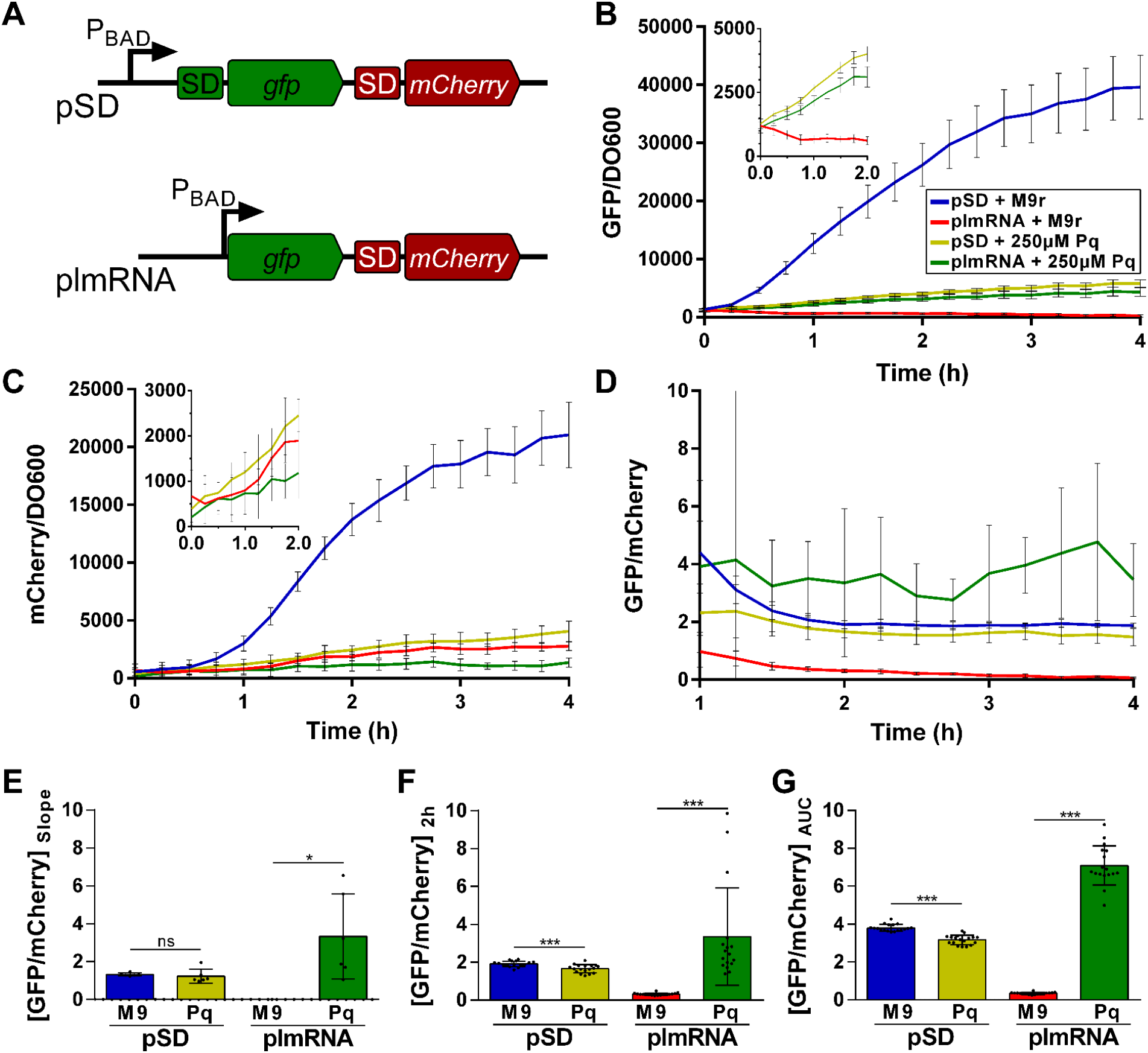
lmRNA is translated more efficiently under oxidative stress. A) Scheme of leader (pSD) or leaderless (plmRNA) reporters used in B, C, D, E, F and G. The biscistronic reporter system codes superfolder GFP (*gfp*) and mCherry (*mCherry*), downstream of the P_BAD_ promoter. B) and C) Kinetics of fluorescence normalized by OD_600_ for GFP (B) and mCherry (C). D) Kinetic of the ratio of fluorescence intensities between GFP and mCherry (GFP/mCherry) for pSD and plmRNA in control (M9) and oxidative stress (250 μM Paraquat) conditions. E) Ratio between the slopes of the linear segment of the GFP/DO600 (0,5-2,0h) and mCherry/DO600 (1-2h) kinetics, under control and stress conditions. F) GFP/mCherry ratio at 2 h after transcription induction, under control and stress conditions. G) Area under the curve (AUC) of the graph in figure 2B between 2 and 4 h after transcription induction, under control and stress condition. Statistical analyses: unpaired two tailed t test with Welch’s correction. ****p<0,0001, ***p<0,001, *p<0,05.

When comparing fluorescence intensity kinetics of both reporters, we observed that translational levels of GFP and mCherry encoded in pSD decrease drastically under oxidative stress (Figure 2BC). In contrast, synthesis of GFP encoded in the leaderless transcript increased under oxidative stress (Figure 2B), while the levels of mCherry (that remains being translated by the canonical mechanism) show a small decrease (Figure 2C). Together, these results indicate that a strong inhibition of canonical initiation of translation occurs under oxidative stress, while efficiency of the translation of lmRNA increases with respect to control condition. As a result, we concluded that *gfp* in an lmRNA context is translated more efficiently than in a canonical context under oxidative stress. This interpretation derives from comparing the ratio between the speed of GFP and mCherry production (Figure 2E). As a plot of the ratio between the values of two straight lines gives an asymptote that tends to the ratio of the slopes of both lines, a similar trend was observed by analyzing the ratio between the levels of GFP and mCherry after 2 hrs of induction (Figure 2DF) or a ratio between the areas under the curves in the linear region of slope ~0 of a GFP/mCherry vs time plot (Figure 2DG). All these approaches indicated an estimated 10 to 20 fold increase for the efficiency of translation of leaderless reporter with respect to the canonical reporter under oxidative stress induced by 250 μM paraquat. pSD reporter does not show relevant changes of the GFP/mCherry ratio because translation efficiency is altered to a similar extent for both *gfp* and *mCherry* when it depends on canonical initiation. As the three methods give similar conclusions, further results are only expressed as the ratio of fluorescence at 2 hrs. To confirm this result does not derive from partial degradation of the reporter mRNAs, qRT-PCR was performed with probes targeted to 3 regions of the bicistronic reporter. As shown in supplementary figure S1, there are small changes on the levels of each region as a consequence of oxidative stress. Under oxidative stress, these changes of mRNA levels would produce a small increase of *gfp* mRNA for the SD reporter and a small increase of *mCherry* mRNA for the lmRNA reporter. Thus, changes of GFP/mCherry fluorescence does not derive form partial mRNA degradation. If anything, this result suggests that activation of lmRNA translation is stronger than interpreted exclusively from fluorescence changes. To further confirm that the GFP/mCherry method is useful to study alterations in lmRNA translation, we repeated this experiment inducing stress with kasugamycin, an antibiotic that is known to induce lmRNA translation (Kaberdina et al., 2009; Schluenzen et al., 2006; Schuwirth et al., 2006). Kasugamycin induced a similar increase in the GFP/mCherry fluorescence ratio for the plmRNA reporter (Supplementary Figure S2) as observed with paraquat (Figure 2), further supporting our conclusion. Thus, *in toto*, our results strongly suggest that 250 μM paraquat induce the translation of lmRNAs. To our knowledge, this is the first report of increased lmRNA translation in conditions that do not derive from antibiotics stress (Kaberdina et al., 2009) or artificial deletion (Landwehr et al., 2021) or overexpression of genes (Vesper et al., 2011).

### MazEF toxin and antitoxin are not involved in the oxidative stress activation of lmRNA translation

Although it has been observed that lmRNAs have a higher affinity for whole ribosomes (70S) than for the minor subunit (30S) (O’Donnell & Janssen, 2002), today there is no clarity on the mechanism responsible for initiation of their translation. The formation of specialized ribosomes, responsible for the preferential translation of mRNA under certain stress conditions has been proposed. This process would be mediated by the MazEF toxin-antitoxin system, which would be responsible for partially digesting the 3’ end of the 16S rRNA of the minor ribosome subunit (30S) and/or complete ribosomes (70S). This mechanism would generate ribosomes lacking 43 nucleotides (70S^Δ43^) that contain the aSD sequence, causing a reduction in the affinity for mRNAs with a leader region, but maintaining translation of leaderless transcripts (Temmel et al., 2016; Vesper et al., 2011).

To evaluate whether the translation changes observed for the plmRNA reporter under oxidative stress depend on the action of MazEF, a mutant of the *mazEF* operon was constructed by homologous recombination with PCR products (Datsenko & Wanner, 2000). From the analysis of the translational kinetics using wild type *E. coli* K-12 MG1655 (K12) and mutant *E. coli* K-12 MG1655 Δ*mazEF*∷FRT (Δ*mazEF*), we observed that deletion of the MazEF toxin-antitoxin system does not prevent the strong increase in translation of *gfp* from the plmRNA reporter under oxidative stress (Figure 3A).

**Figure 3.**
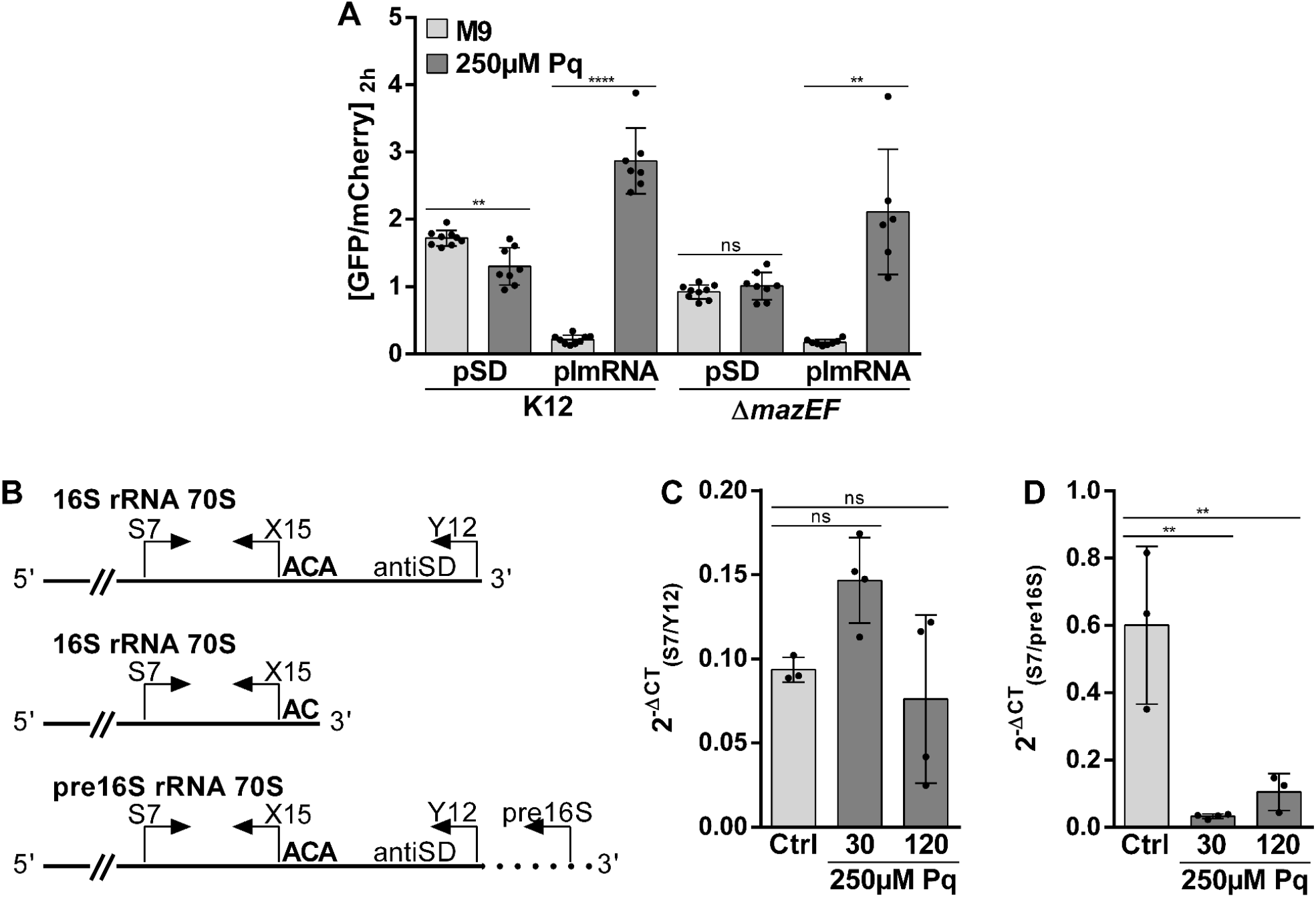
MazF and the cleavage of 16S 3’ extreme are not required to induce translation under oxidative stress. A) GFP/mCherry fluorescence ratio for wild-type and Δ*mazEF*∷FRT (Δ*mazEF*) *E. coli* strains carrying pSD and plmRNA constructions at 2 h after transcription induction in control and stress condition. B) PCR strategy used to analyse the integrity of 16S rRNA 3’ extreme. C) and D) Integrity analysis of 16S rRNA (C) and pre 16S rRNA (D) from wild-type *E. coli* under control condition and 30 or 120 min after oxidative stress induction. Statistical analyses: unpaired two-tailed t test for A. One-way ANOVA and Dunnett’s multiple comparisons test for C and D. ****p<0,0001, **p<0,01, *p<0,05.

To further verify whether MazF or another endoribonuclease are processing the 16S rRNA under oxidative stress induced by paraquat, the integrity of its 3’ end was analyzed by real-time RT-PCR. To this end, primers that hybridize at both sides of the proposed ACA cleavage site and indicate the levels of the non-cleaved 16S rRNA were used (primers S7 to Y12 in Figure 3B) (Vesper et al., 2011). A second pair of primers (S7 and X15) hybridizing a region conserved in all 16S rRNA species independent of their processing were used as an internal control of 16S rRNA levels. When analyzing the wild type strain, no significant changes were observed in the relative levels of the complete 3’ end of the 16S rRNA after induction of oxidative stress with 250 μM of paraquat. This suggests the absence of digestion at the 3’ end of the 16S rRNA under oxidative stress (Figure 3C). Confirming these results, we were unable to detect the expected 43 nucleotide fragment by Northern blot analysis using the same probes that have been previously reported (Vesper et al., 2011) (data not shown).

Although in contradiction to the MazF model for lmRNA translation activation, these results coincide with reports from other researchers that have suggested that the MazF endoribonuclease would not generate 70S^Δ43^ ribosomes. Instead, these researchers found that activation of MazF non-selectively degrades immature rRNA and mRNA (Culviner & Laub, 2018; Mets et al., 2017). In accordance with those reports, when analyzing the levels of the 16S rRNA precursor using a primer that hybridizes to a region of the immature rRNA (downstream of the mature 3’ extreme, Figure 3B), we observed that the levels of 16S pre - rRNA decrease dramatically after the induction of oxidative stress (Figure 3D). All together, these results suggest that in accordance with what was published by Mets *et al*., 2017 and Culviner & Laub, 2018, in *E. coli* K-12 MG1655, 70S^Δ43^ ribosomes are not generated under oxidative stress. The results are also congruent with a decreased abundance of ribosomal RNA precursors that may derive from either their degradation or decreased transcription.

### Oxidative stress increases the translational efficiency of lmRNAs naturally found in *E. coli*

Our results strongly supports that translation of lmRNA is activated under oxidative stress (Figure 2) in a mechanism that probably involves some changes in the composition of the ribosomal particles (Figure 1), but is independent of the processing of the 16S rRNA 3’ extreme (Figure 3). Nevertheless, all these analyzes depend on experiments carried out using the plasmid plmRNA, which generates an artificial leaderless mRNA encoding the heterologous genes *gfp* and *mCherry*. However, leaderless transcripts can also be found in nature. For example, as a product of alternative promoters in regulatory genes from phages λ and P2, transposon Tn1721, or *virF* in *Shigella* spp. (Cortes et al., 2013; Di Martino et al., 2016; Resch et al., 1995). The first approaches to understand the mechanism of leaderless translation were carried out with the *cI* gene of phage λ. This gene has two described promoters, P_RE_ (promoter for repressor establishment) and P_RM_ (promoter for repressor maintenance). Transcription from P_RE_ generates a transcript with a 5’ untranslated region of 403 nucleotides, while transcription beginning at P_RM_ produces a leaderless mRNA, whose 5’ end corresponds to the AUG start codon (Schmeissner et al., 1980; Walz et al., 1976). To evaluate changes in the translation efficiency of the leaderless *cI* under oxidative stress, a translation fusion of the first 100 *cI* codons (from P_RM_) to *gfp* was constructed in the vector plmRNA, forming reporter *cI-gfp* (Figure 4A). As expected, we observed an increase in GFP/mCherry ratio under oxidative stress (Figure 4C). Although activation of lmRNA translation is clear, this enhancement is more discreet than what we have observed using the plmRNA reporter (Figure 2). We are currently unable to explain this difference, but it is possible that it derives from differences between the sequence of the first codons of both reporters of lmRNA translation, as previous reports have shown idiosyncratic effects of diverse lmRNA reporter sequences (Beck et al., 2016; Shell et al., 2015; Udagawa et al., 2004).

**Figure 4.**
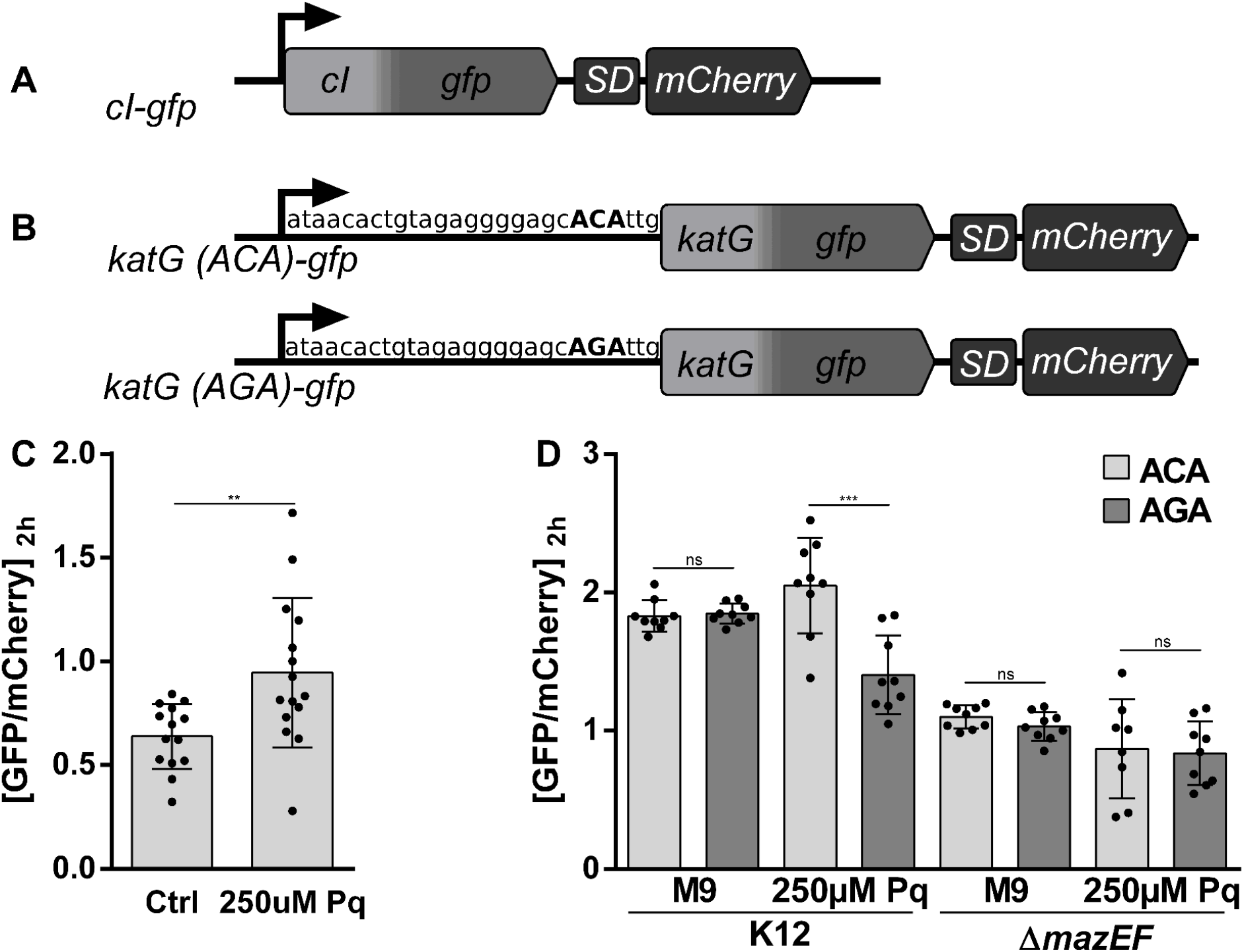
Oxidative stress increases the efficiency of translation of genes naturally encoded as leaderless transcripts. A) Scheme of the translational fusion of the first 100 codons of leaderless *cI* transcript (from P_RM_ promoter) to the *gfp-mCherry* reporter system. B) Scheme of the translational fusion between the 5’UTR and the first 100 codons of *katG* transcript to *gfp-mCherry* reporter system. *katG(ACA)-gfp* includes the wild-type 5’UTR while *katg(AGA)-gfp* includes a point mutation in sequence ACA, recognized by MazF. C) GFP/mCherry fluorescence ratio of *cI-gfp* fusion at 2 h after transcription induction, in control and stress conditions. D) GFP/mCherry fluorescence ratio of *katG-gfp* fusion at 2 h after transcription induction, in control and stress conditions in the WT and Δ*mazEF* strains. Statistical analyses: unpaired two-tailed t test with Welch’s correction for C. Unpaired two-tailed t test for D. ***p<0,001, **p<0,01.

In addition to the synthesis of lmRNA by transcription from promoters located near to the initiation codon, some authors have proposed that lmRNAs may be also generated through cleavage by MazF of canonical mRNAs (Vesper et al., 2011). Possible MazF targets with their respective ACA cleavage sites have been described in *E. coli* strains that overexpress *mazF* (Sauert et al., 2016). Four of these genes belong to the SoxRS and OxyR regulons and thus, are potentially involved in the response to oxidative stress by paraquat: *fpr*, *katG*, *nepI* and *zwf* (Seo et al., 2015). Since the transcription initiation site of *katG* has been described (Tartaglia et al., 1989), this gene was chosen to study whether MazF may be involved in the production of lmRNAs under oxidative stress. To this end, a translation fusion between the 5’ extreme of *katG* (including the 5’ UTR and the first 100 codons of the gene) and the *gfp* reporter was constructed. Furthermore, a point mutation was generated in the ACA cleavage site replacing it with AGA, to obtain a transcript that is not processable by MazF (Figure 4B). When analyzing *gfp* translation in the wild type strain, we observed no differences between the *katG*(ACA)-*gfp* and *katG*(AGA)-*gfp* reporters under control conditions (Figure 4D). However, under oxidative stress induced by 250 μM of paraquat, we observed a higher GFP/mCherry ratio for the reporter that can be processed by MazF (ACA) (Figure 4D). When repeating these experiments in a strain lacking the toxin-antitoxin system, we did not observe any differences between the reporters, both under control and oxidative stress conditions (Figure 4D). Since we previously observed that MazF does not play a role in the increased translation of lmRNA under oxidative stress (Figure 3A), these latest data suggest that MazF would participate in the processing of the *katG* transcript, increasing its translational efficiency under oxidative stress. As reporters differ only in the cleavage site described for MazF, it can be proposed that the increased translation of *katG*(ACA)-*gfp* is a consequence of the more efficient translation of at least a fraction of the mRNA that is processed into lmRNA. This result is similar to that recently described for the *grcA* mRNA (MazF target), whose translation increases in a part of the bacterial population when MazF is overexpressed (Nikolic et al., 2017). Nevertheless, care should be taken in considering that induction of lmRNA translation under stress conditions in the WT strain is modest, an effect that may derive from further digestion of transcripts cleaved by MazF (Culviner & Laub, 2018).

### (p)ppGpp accumulation inhibits translation of canonical mRNAs under oxidative stress

Our results indicate that under stress conditions, a decrease in ribosomal RNA precursors is observed (Figure 3D). Such decrease may be a consequence of the accumulation of guanosine tetra- or penta-phosphate ((p)ppGpp) (Sanchez-Vazquez et al., 2019). This is a nucleotide alarmone produced when *E. coli* or other bacteria are exposed to decreased amino acids availability or several other conditions, including oxidative stress (Chang et al., 2002). It has been determined that this molecule is capable of interacting with around 30 proteins involved in different processes, including some involved in ribosome biogenesis and translation. As a consequence, (p)ppGpp inhibits transcription and translation of several genes (Zhang et al., 2018). Although it is well known that transcription of some genes is enhanced by (p)ppGpp (Sanchez-Vazquez et al., 2019), we are unaware of any report regarding its effects on lmRNA translation. To determine whether this metabolite plays a role in the regulation of leaderless translation, we analyzed the efficiency of GFP production from pSD and plmRNA reporters in a mutant lacking the *relA* and *spoT* genes (Δ*relA*∷FRT Δ*spoT*∷FRT or Δ*relA*Δ*spoT*), responsible for the synthesis of (p)ppGpp in *E. coli* (46). As mutants that cannot produce (p)ppGpp are unable to replicate in culture media devoid of most amino acids (Potrykus et al., 2011), we initially tried to perform the experiments in M9 media supplemented with triptone, a casein digest that contains all amino acids. Unexpectedly, in this culture medium we did not observe lmRNA translation in either the WT or the Δ*relA*Δ*spoT* strains, both under control or oxidative stress conditions (Data not shown). To overcome this limitation, WT and Δ*relA*Δ*spoT* strains were cultured in M9 media supplemented with tryptone. Before performing the experiment, cultures were cleaned in M9 media and resuspended in M9 media with branched amino acids where they were used to test the production of GFP and mCherry. Using this protocol, we observed that lmRNA translation is activated in the WT strain under oxidative stress, replicating our previous observations. Nevertheless, the Δ*relA*Δ*spoT* strain was unable to translate the lmRNA *gfp* reporter under any conditions. If we instead resuspended the cell in M9 media containing tryptone, the lmRNA reporter was not translated in any of the conditions or strains (Figure 5). Interestingly, deletion of the genes involved in (p)ppGpp synthesis also prevented the inhibition of canonical GFP and mCherry translation under oxidative stress (Figure 5). Altogether, these results show that lmRNA cannot be translated in strains impeded to produce sufficient levels of (p)ppGpp either due to mutation of *relA* and *spoT* or because of a high amino acids availability. Thus, our work strongly suggests that in addition to inhibiting canonical translation, (p)ppGpp is required for the activation of lmRNA translation at least under oxidative stress. Deletion of only *relA* was not enough to prevent lmRNA translation under oxidative stress, although it did affect translation of canonical mRNAs (Supplementary Figure S3).

**Figure 5.**
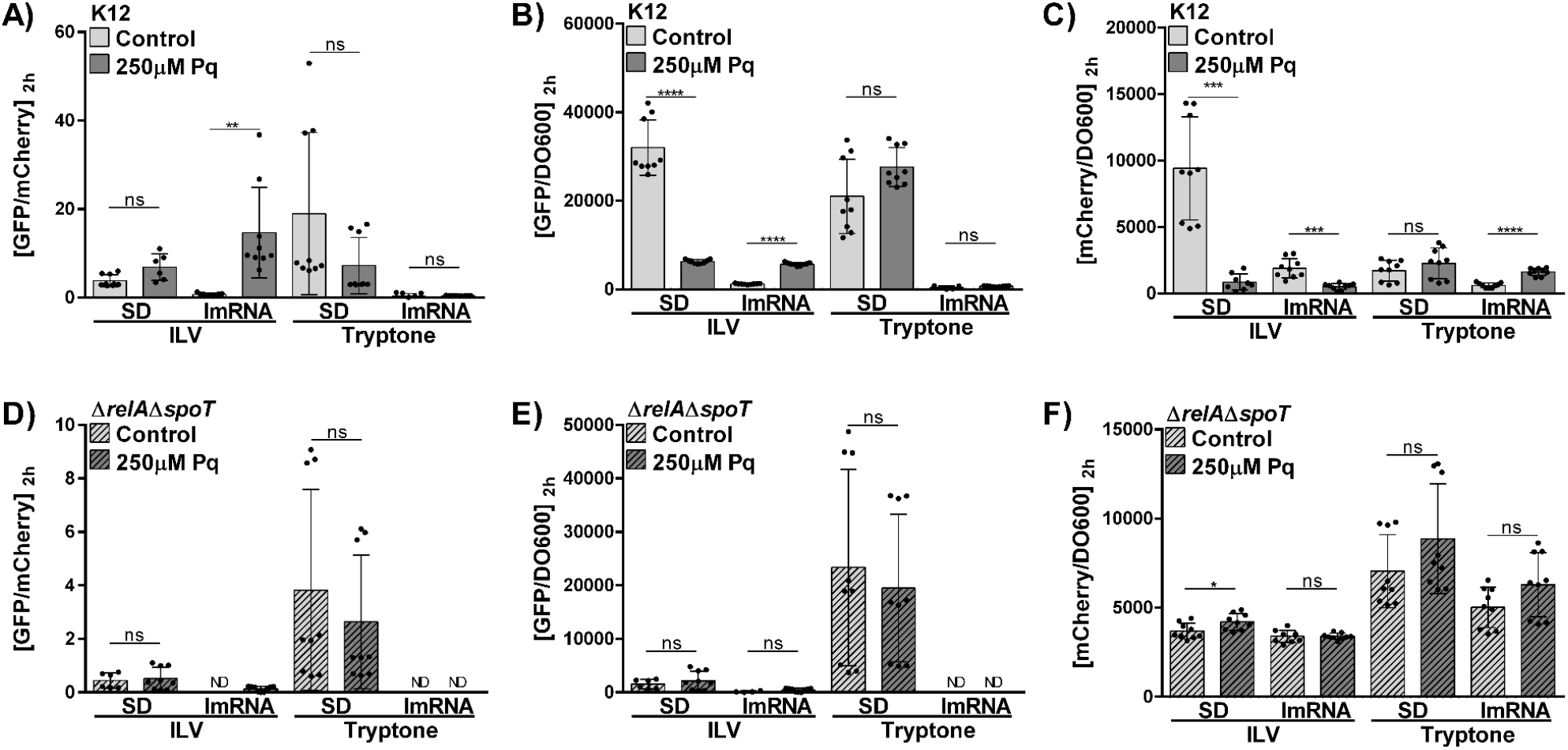
The induction of lmRNA translation under oxidative stress depends on the synthesis of (p)ppGpp and the availability of amino acids. GFP/mCherry, GFP/DO600 and mCherry/DO600 ratios for WT (K12; A, B and C) and Δ*relA*∷FRT Δ*spoT*∷FRT (Δ*relA*Δ*spoT*; D, E and F) *E. coli* strains carrying the pSD and plmRNA constructs, at 2 h after transcription induction in control (light gray) and oxidative stress conditions (dark gray). The M9 culture medium was supplemented with low concentrations of branched amino acids (ILV 50μg/mL each) or high concentration of all amino acids (Tryptone 0.5%). Note that experiments in this figure were performed with three independent clones of the Δ*relA*Δ*spoT* strain, explaining in part the higher dispersion of data. ND= Not detected. Statistical analyses: unpaired two tailed t test with Welch’s correction. ****p<0,0001, ***p<0,001, **p<0,01, *p<0,05.

Part of the (p)ppGpp response is mediated by Lon protease activation that leads to degradation of ribosomal proteins (Kuroda, 2001). However, *lon* deletion does not prevent *gfp* expression from the lmRNA reporter (Supplementary Figure S3) indicating that lmRNA translation does not require the Lon dependent degradation of ribosomal proteins.

## Discussion

Because initiation is usually the slowest step of translation, regulation of mRNA translation usually depends on alterations of this step. This is usually achieved by changes to the accessibility of the RBS, ultimately controlling binding of the 30S subunit (Duval et al., 2015). In addition to this regulation of translation of particular genes, global inhibition of translation has been observed as a response to several hostile conditions such as the stationary phase of bacterial cultures (Kato et al., 2010), when Mg^+2^ is limiting (Pontes et al., 2016) or as a response to cold shock (Vila-Sanjurjo et al., 2004). This decrease in translation usually depends on the inhibition of ribosomes, but in some cases changes in the accessibility of the RBS also plays a role. One of the best described mechanisms for global inhibition of bacterial translation is the early response to cold shock in *E. coli*. As temperature decreases, mRNA folds, preventing access of the ribosome to correct translation initiation sites. Thus, translation must be globally repressed, preventing initiation at incorrect sites. As a response, most *E. coli* ribosomes enter a hibernation state induced by binding of YfiA protein to the P and A sites of complete ribosomes (70S). This YfiA-70S complex is unable to catalyze translation (Vila-Sanjurjo et al., 2004). Nevertheless, in order to confront the cold shock, it is not enough to block mis-translation of folded mRNA. Bacteria must additionally enhance translation of genes that may aid its ability to overcome the stress conditions. It has been observed that the reduced number of functional ribosomes selectively translate a minor subset of the available mRNAs that code for RNA chaperons and other proteins that allow a better adaptation to cold conditions. All these mRNA fold under cold conditions to a structure where the RBS and enhancer sequences remain exposed to binding by the 30S allowing translation initiation (Vila-Sanjurjo et al., 2004). Thus, the early cold shock response is characterized by a global inhibition of translation together with the induction of translation of a subset of genes required for cold shock response.

The response to oxidative stress seems to follow a similar behavior to what has been observed under cold shock, although mechanisms appear to be different. Several authors have observed a global inhibition of translation (Kojima et al., 2007; Nishiyama et al., 2004; Shenton et al., 2006; Zhong et al., 2015; M. Zhu & Dai, 2019). Some of them have indicated that this inhibition would derive from massive tRNA degradation (Zhong et al., 2015; M. Zhu & Dai, 2019). Nevertheless, it seems that this would be a strain dependent behavior, as strain K-12 MG1655 used for this report does not show such massive decrease in tRNA content (Leiva et al., 2020). Here we have observed that (p)ppGpp production may be responsible for at least part of such inhibition. Perhaps, this may derive from the ability of (p)ppGpp to bind to some translation factors using the GTP binding sites (Kanjee et al., 2012). Interestingly, this is the case of IF2 (Mitkevich et al., 2010) that is required for canonical, but not for lmRNA translation (Udagawa et al., 2004). Additional inhibition may derive from direct oxidation of proteins and RNA involved in the translation process (Katz & Orellana, 2012; Liu et al., 2012; Willi et al., 2018). Independent of the mechanism underlying global translation inhibition, bacteria will need to preferentially translate genes that allow adaptation to oxidative conditions. Otherwise, the effects of transcriptional control on bacterial adaptation would be limited by a slow translation apparatus. Here we have observed that translation of lmRNA is activated under oxidative stress conditions, apparently by the same (p)ppGpp that inhibits canonical translation. This activation may be an indirect consequence of the increased availability of ribosomes that show decreased translation of canonical mRNAs when translation is inhibited by (p)ppGpp. Also, high levels of aminoacylated initiator tRNA, which probably increase as a consequence of canonical translation inhibition, have been shown to enhance lmRNA translation *in vitro* and could also be involved in the regulatory process (Udagawa et al., 2004). Recently, the ATPase YchF of *E. coli* has been shown to bind the ribosomes and apparently inhibit lmRNA translation (Landwehr et al., 2021). Thus, it is possible that (p)ppGpp may inhibit YchF, although it is also possible that YchF somehow prevents (p)ppGpp production or that it controls lmRNA translation by a (p)ppGpp independent mechanism.

Understanding the mechanisms involved in (p)ppGpp activation of lmRNA translation will require further investigation. Nevertheless, the apparent ability of (p)ppGpp to simultaneously inhibit canonical translation and enhance lmRNA translation could allow for the selective translation of a subset of the transcriptome under oxidative stress. Additional experiments will be also required to determine this subset of lmRNAs, allowing a better understanding of the translational control for the adaptation to oxidative stress. If confirmed, these results suggest that bacteria may confront diverse hostile conditions through a common strategy of translation regulation, that is, a global inhibition of translation together with a concomitant activation of the translation of particular genes.

## Materials and Methods

A brief description of the main methods used in this work is provided here. A detailed description can be found at the supplementary methods section.

### Strains and culture media

All experiments performed in this work used wild-type (WT) *E. coli* K-12 MG1655 strain. Strains were cultured in either lysogeny broth (LB) media (1% tryptone, 0.5% yeast extract, and 0.5% NaCl), or M9 media (47.7 mM Na_2_HPO_4_, 22.0 mM KH_2_PO_4_, 8.6 mM NaCl, 18.7 mM NH_4_Cl, 2 mM MgSO_4_, 0.1 mM CaCl_2_, and 0.4% Glycerol). When indicated, branched amino acids (isoleucine, leucine, and valine 50 μg/ml each), 0.1% tryptone, 0.4% arabinose 0.4%, ampicillin (100 μg/ml) or 250 μM paraquat were added to the culture media.

### Analysis of polysomes

The detailed protocol for analysis of polysomes has been described by Qin and Kurt (2013). In brief, 50 ml of M9 media supplemented with branched amino acids were inoculated with saturated overnight culture in M9 media supplemented with tryptone and grown at 37°C in an orbital shaker. When indicated, the media also contained 250 μM paraquat. When bacteria reached mid-log phase (OD 600 ~0.4–0.6), cells were collected, lized and prepared for ultracentrifugation as indicated in the supplementary methods. Samples were then loaded on a 10%-40% sucrose gradient and centrifuged in a SW41 rotor at 35000 RPM for 3 hours. Finally, samples were analyzed using an in-line UV detector (ISCO/Brandel system).

### Translation Efficiency Analyses

50 μl aliquot of mid-log phase cultures were diluted in a 96-well optical-bottom plate with 150 μl fresh M9 media supplemented with branched amino acids (50 μg/ml each) and arabinose (0.4% final concentration). When indicated, culture media contained paraquat (250 μM final concentration). Also 0,5% tryptone was used instead of branched amino acids in some experiments. Plates were further shaken at 37°C. OD 600 and fluorescence intensity of GFP (Ex. 480 ± 4.5 nm, Em. 515 ± 10 nm) and mCherry (Ex. 555 ± 4.5 nm, Em. 600 ± 10 nm) were measured in a microplate reader (Infinite M200 PRO, Tecan). In all experiments, strains transformed with pBAD30 (parental plasmid of pBAD30SFIT) were used to subtract the fluorescence background. A detailed description of the protocol can be found at the supplementary methods section.

## Author contribution

The project was originally designed by MI, AK and OO. All experiments were performed by LL. AK, MI and OO supervised experiments and data interpretation. LL and AK wrote the paper with contributions from all authors. All authors reviewed and approved the final version of the manuscript.

## References

Beck, H. J., Fleming, I. M. C., & Janssen, G. R. (2016). 5’-Terminal AUGs in *Escherichia coli* mRNAs with Shine-Dalgarno Sequences: Identification and Analysis of Their Roles in Non-Canonical Translation Initiation. PLOS ONE, 11(7), e0160144. https://doi.org/10.1371/journal.pone.0160144

Chang, D.-E., Smalley, D. J., & Conway, T. (2002). Gene expression profiling of *Escherichia coli* growth transitions: An expanded stringent response model. Molecular Microbiology, 45(2), 289–306. https://doi.org/10.1046/j.1365-2958.2002.03001.x

Chiang, S. M., & Schellhorn, H. E. (2012). Regulators of oxidative stress response genes in *Escherichia coli* and their functional conservation in bacteria. Archives of Biochemistry and Biophysics, 525(2), 161–169. https://doi.org/10.1016/j.abb.2012.02.007

Cortes, T., Schubert, O. T., Rose, G., Arnvig, K. B., Comas, I., Aebersold, R., & Young, D. B. (2013). Genome-wide Mapping of Transcriptional Start Sites Defines an Extensive Leaderless Transcriptome in *Mycobacterium tuberculosis*. Cell Reports, 5(4), 1121–1131. https://doi.org/10.1016/j.celrep.2013.10.031

Culviner, P. H., & Laub, M. T. (2018). Global Analysis of the *E. coli* Toxin MazF Reveals Widespread Cleavage of mRNA and the Inhibition of rRNA Maturation and Ribosome Biogenesis. Molecular Cell, 70(5), 868–880.e10. https://doi.org/10.1016/j.molcel.2018.04.026

Datsenko, K. A., & Wanner, B. L. (2000). One-step inactivation of chromosomal genes in Escherichia coli K-12 using PCR products. Proceedings of the National Academy of Sciences, 97(12), 6640–6645. https://doi.org/10.1073/pnas.120163297

de Groot, A., Roche, D., Fernandez, B., Ludanyi, M., Cruveiller, S., Pignol, D., Vallenet, D., Armengaud, J., & Blanchard, L. (2014). RNA Sequencing and Proteogenomics Reveal the Importance of Leaderless mRNAs in the Radiation-Tolerant Bacterium Deinococcus deserti. Genome Biology and Evolution, 6(4), 932–948. https://doi.org/10.1093/gbe/evu069

Delvillani, F., Papiani, G., Dehò, G., & Briani, F. (2011). S1 ribosomal protein and the interplay between translation and mRNA decay. Nucleic Acids Research, 39(17), 7702–7715. https://doi.org/10.1093/nar/gkr417

Di Martino, M. L., Romilly, C., Wagner, E. G. H., Colonna, B., & Prosseda, G. (2016). One Gene and Two Proteins: A Leaderless mRNA Supports the Translation of a Shorter Form of the *Shigella* VirF Regulator. MBio, 7(6), e01860–16. https://doi.org/10.1128/mBio.01860-16

Duval, M., Simonetti, A., Caldelari, I., & Marzi, S. (2015). Multiple ways to regulate translation initiation in bacteria: Mechanisms, regulatory circuits, dynamics. Biochimie, 114, 18–29. https://doi.org/10.1016/j.biochi.2015.03.007

Fasnacht, M., & Polacek, N. (2021). Oxidative Stress in Bacteria and the Central Dogma of Molecular Biology. Frontiers in Molecular Biosciences, 8, 671037. https://doi.org/10.3389/fmolb.2021.671037

Gualerzi, C. O., & Pon, C. L. (2015). Initiation of mRNA translation in bacteria: Structural and dynamic aspects. Cellular and Molecular Life Sciences, 72(22), 4341–4367. https://doi.org/10.1007/s00018-015-2010-3

Hauryliuk, V., Atkinson, G. C., Murakami, K. S., Tenson, T., & Gerdes, K. (2015). Recent functional insights into the role of (p)ppGpp in bacterial physiology. Nature Reviews Microbiology, 13(5), 298–309. https://doi.org/10.1038/nrmicro3448

Hwang, S., Lee, N., Jeong, Y., Lee, Y., Kim, W., Cho, S., Palsson, B. O., & Cho, B.-K. (2019). Primary transcriptome and translatome analysis determines transcriptional and translational regulatory elements encoded in the *Streptomyces clavuligerus* genome. Nucleic Acids Research, 47(12), 6114–6129. https://doi.org/10.1093/nar/gkz471

Imlay, J. A. (2013). The molecular mechanisms and physiological consequences of oxidative stress: Lessons from a model bacterium. Nature Reviews Microbiology, 11(7), 443–454. https://doi.org/10.1038/nrmicro3032

Imlay, J. A. (2019). Where in the world do bacteria experience oxidative stress?: Oxidative stress in natural environments. Environmental Microbiology, 21(2), 521–530. https://doi.org/10.1111/1462-2920.14445

Kaberdina, A. C., Szaflarski, W., Nierhaus, K. H., & Moll, I. (2009). An Unexpected Type of Ribosomes Induced by Kasugamycin: A Look into Ancestral Times of Protein Synthesis? Molecular Cell, 33(2), 227–236. https://doi.org/10.1016/j.molcel.2008.12.014

Kaminishi, T., Wilson, D. N., Takemoto, C., Harms, J. M., Kawazoe, M., Schluenzen, F., Hanawa-Suetsugu, K., Shirouzu, M., Fucini, P., & Yokoyama, S. (2007). A Snapshot of the 30S Ribosomal Subunit Capturing mRNA via the Shine-Dalgarno Interaction. Structure, 15(3), 289–297. https://doi.org/10.1016/j.str.2006.12.008

Kanjee, U., Ogata, K., & Houry, W. A. (2012). Direct binding targets of the stringent response alarmone (p)ppGpp: Protein targets of ppGpp. Molecular Microbiology, 85(6), 1029–1043. https://doi.org/10.1111/j.1365-2958.2012.08177.x

Kataria, R., & Ruhal, R. (2014). Microbiological Metabolism Under Chemical Stress. In Microbial Biodegradation and Bioremediation (pp. 497–509). Elsevier. https://doi.org/10.1016/B978-0-12-800021-2.00021-2

Kato, T., Yoshida, H., Miyata, T., Maki, Y., Wada, A., & Namba, K. (2010). Structure of the 100S Ribosome in the Hibernation Stage Revealed by Electron Cryomicroscopy. Structure, 18(6), 719–724. https://doi.org/10.1016/j.str.2010.02.017

Katz, A., & Orellana, O. (2012). Protein Synthesis and the Stress Response. In M. Biyani (Ed.), Cell-Free Protein Synthesis. InTech. https://doi.org/10.5772/50311

Kojima, K., Oshita, M., Nanjo, Y., Kasai, K., Tozawa, Y., Hayashi, H., & Nishiyama, Y. (2007). Oxidation of elongation factor G inhibits the synthesis of the D1 protein of photosystem II. Molecular Microbiology, 65(4), 936–947. https://doi.org/10.1111/j.1365-2958.2007.05836.x

Komarova, A. V., Tchufistova, L. S., Supina, E. V., & Boni, I. V. (2002). Protein S1 counteracts the inhibitory effect of the extended Shine-Dalgarno sequence on translation. RNA, 8(9), 1137–1147. https://doi.org/10.1017/S1355838202029990

Kröger, C., Dillon, S. C., Cameron, A. D. S., Papenfort, K., Sivasankaran, S. K., Hokamp, K., Chao, Y., Sittka, A., Hébrard, M., Händler, K., Colgan, A., Leekitcharoenphon, P., Langridge, G. C., Lohan, A. J., Loftus, B., Lucchini, S., Ussery, D. W., Dorman, C. J., Thomson, N. R., … Hinton, J. C. D. (2012). The transcriptional landscape and small RNAs of *Salmonella enterica* serovar Typhimurium. Proceedings of the National Academy of Sciences, 109(20), E1277–E1286. https://doi.org/10.1073/pnas.1201061109

Kuroda, A. (2001). Role of Inorganic Polyphosphate in Promoting Ribosomal Protein Degradation by the Lon Protease in *E. coli*. Science, 293(5530), 705–708. https://doi.org/10.1126/science.1061315

Landwehr, V., Milanov, M., Angebauer, L., Hong, J., Jüngert, G., Hiersemenzel, A., Siebler, A., Schmit, F., Öztürk, Y., Dannenmaier, S., Drepper, F., Warscheid, B., & Koch, H.-G. (2021). The Universally Conserved ATPase YchF Regulates Translation of Leaderless mRNA in Response to Stress Conditions. Frontiers in Molecular Biosciences, 8, 643696. https://doi.org/10.3389/fmolb.2021.643696

Langevin, S. A., Bent, Z. W., Solberg, O. D., Curtis, D. J., Lane, P. D., Williams, K. P., Schoeniger, J. S., Sinha, A., Lane, T. W., & Branda, S. S. (2013). Peregrine: A rapid and unbiased method to produce strand-specific RNA-Seq libraries from small quantities of starting material. RNA Biology, 10(4), 502–515. https://doi.org/10.4161/rna.24284

Laursen, B. S., Sørensen, H. P., Mortensen, K. K., & Sperling-Petersen, H. U. (2005). Initiation of Protein Synthesis in Bacteria. Microbiology and Molecular Biology Reviews, 69(1), 101–123. https://doi.org/10.1128/MMBR.69.1.101-123.2005

Leiva, L. E., Pincheira, A., Elgamal, S., Kienast, S. D., Bravo, V., Leufken, J., Gutiérrez, D., Leidel, S. A., Ibba, M., & Katz, A. (2020). Modulation of *Escherichia coli* Translation by the Specific Inactivation of tRNA^Gly^ Under Oxidative Stress. Frontiers in Genetics, 11, 856. https://doi.org/10.3389/fgene.2020.00856

Ling, J., & Soll, D. (2010). Severe oxidative stress induces protein mistranslation through impairment of an aminoacyl-tRNA synthetase editing site. Proceedings of the National Academy of Sciences, 107(9), 4028–4033. https://doi.org/10.1073/pnas.1000315107

Liu, M., Gong, X., Alluri, R. K., Wu, J., Sablo, T., & Li, Z. (2012). Characterization of RNA damage under oxidative stress in *Escherichia coli*. Biological Chemistry, 393(3), 123–132. https://doi.org/10.1515/hsz-2011-0247

Mendoza-Vargas, A., Olvera, L., Olvera, M., Grande, R., Vega-Alvarado, L., Taboada, B., Jimenez-Jacinto, V., Salgado, H., Juárez, K., Contreras-Moreira, B., Huerta, A. M., Collado-Vides, J., & Morett, E. (2009). Genome-Wide Identification of Transcription Start Sites, Promoters and Transcription Factor Binding Sites in E. coli. PLoS ONE, 4(10), e7526. https://doi.org/10.1371/journal.pone.0007526

Mets, T., Lippus, M., Schryer, D., Liiv, A., Kasari, V., Paier, A., Maiväli, Ü., Remme, J., Tenson, T., & Kaldalu, N. (2017). Toxins MazF and MqsR cleave *Escherichia coli* rRNA precursors at multiple sites. RNA Biology, 14(1), 124–135. https://doi.org/10.1080/15476286.2016.1259784

Mitkevich, V. A., Ermakov, A., Kulikova, A. A., Tankov, S., Shyp, V., Soosaar, A., Tenson, T., Makarov, A. A., Ehrenberg, M., & Hauryliuk, V. (2010). Thermodynamic Characterization of ppGpp Binding to EF-G or IF2 and of Initiator tRNA Binding to Free IF2 in the Presence of GDP, GTP, or ppGpp. Journal of Molecular Biology, 402(5), 838–846. https://doi.org/10.1016/j.jmb.2010.08.016

Nguyen, G. T., Green, E. R., & Mecsas, J. (2017). Neutrophils to the ROScue: Mechanisms of NADPH Oxidase Activation and Bacterial Resistance. Frontiers in Cellular and Infection Microbiology, 7, 373. https://doi.org/10.3389/fcimb.2017.00373

Nguyen, T. G., Vargas-Blanco, D. A., Roberts, L. A., & Shell, S. S. (2020). The Impact of Leadered and Leaderless Gene Structures on Translation Efficiency, Transcript Stability, and Predicted Transcription Rates in Mycobacterium smegmatis. Journal of Bacteriology, 202(9), e00746–19. https://doi.org/10.1128/JB.00746-19

Nikolic, N., Didara, Z., & Moll, I. (2017). MazF activation promotes translational heterogeneity of the *grcA* mRNA in *Escherichia coli* populations. PeerJ, 5, e3830. https://doi.org/10.7717/peerj.3830

Nishiyama, Y., Allakhverdiev, S. I., Yamamoto, H., Hayashi, H., & Murata, N. (2004). Singlet Oxygen Inhibits the Repair of Photosystem II by Suppressing the Translation Elongation of the D1 Protein in *Synechocystis* sp. PCC 6803. Biochemistry, 43(35), 11321–11330. https://doi.org/10.1021/bi036178q

O’Donnell, S. M., & Janssen, G. R. (2002). Leaderless mRNAs Bind 70S Ribosomes More Strongly than 30S Ribosomal Subunits in *Escherichia coli*. Journal of Bacteriology, 184(23), 6730–6733. https://doi.org/10.1128/JB.184.23.6730-6733.2002

Pontes, M. H., Yeom, J., & Groisman, E. A. (2016). Reducing Ribosome Biosynthesis Promotes Translation during Low Mg 2+ Stress. Molecular Cell, 64(3), 480–492. https://doi.org/10.1016/j.molcel.2016.05.008

Potrykus, K., Murphy, H., Philippe, N., & Cashel, M. (2011). ppGpp is the major source of growth rate control in *E. coli*: PpGpp and growth rate control. Environmental Microbiology, 13(3), 563–575. https://doi.org/10.1111/j.1462-2920.2010.02357.x

Qin, D., & Fredrick, K. (2013). Analysis of Polysomes from Bacteria. In Methods in Enzymology (Vol. 530, pp. 159–172). Elsevier. https://doi.org/10.1016/B978-0-12-420037-1.00008-7

Resch, A., Tedin, K., Graschopf, A., Haggård-Ljungquist, E., & Bläsi, U. (1995). Ternary complex formation on leaderless phage mRNA. FEMS Microbiology Reviews, 17(1–2), 151–157. https://doi.org/10.1111/j.1574-6976.1995.tb00197.x

Rothschild, L. J., & Mancinelli, R. L. (2001). Life in extreme environments. Nature, 409(6823), 1092–1101. https://doi.org/10.1038/35059215

Saito, K., Green, R., & Buskirk, A. R. (2020). Translational initiation in *E. coli* occurs at the correct sites genome-wide in the absence of mRNA-rRNA base-pairing. ELife, 9, e55002. https://doi.org/10.7554/eLife.55002

Sanchez-Vazquez, P., Dewey, C. N., Kitten, N., Ross, W., & Gourse, R. L. (2019). Genome-wide effects on *Escherichia coli* transcription from ppGpp binding to its two sites on RNA polymerase. Proceedings of the National Academy of Sciences, 116(17), 8310–8319. https://doi.org/10.1073/pnas.1819682116

Sauert, M., Wolfinger, M. T., Vesper, O., Müller, C., Byrgazov, K., & Moll, I. (2016). The MazF-regulon: A toolbox for the post-transcriptional stress response in *Escherichia coli*. Nucleic Acids Research, 44(14), 6660–6675. https://doi.org/10.1093/nar/gkw115

Schluenzen, F., Takemoto, C., Wilson, D. N., Kaminishi, T., Harms, J. M., Hanawa-Suetsugu, K., Szaflarski, W., Kawazoe, M., Shirouzu, M., Shirouzo, M., Nierhaus, K. H., Yokoyama, S., & Fucini, P. (2006). The antibiotic kasugamycin mimics mRNA nucleotides to destabilize tRNA binding and inhibit canonical translation initiation. Nature Structural & Molecular Biology, 13(10), 871–878. https://doi.org/10.1038/nsmb1145

Schmeing, T. M., & Ramakrishnan, V. (2009). What recent ribosome structures have revealed about the mechanism of translation. Nature, 461(7268), 1234–1242. https://doi.org/10.1038/nature08403

Schmeissner, U., Court, D., Shimatake, H., & Rosenberg, M. (1980). Promoter for the establishment of repressor synthesis in bacteriophage lambda. Proceedings of the National Academy of Sciences, 77(6), 3191–3195. https://doi.org/10.1073/pnas.77.6.3191

Schuwirth, B. S., Day, J. M., Hau, C. W., Janssen, G. R., Dahlberg, A. E., Cate, J. H. D., & Vila-Sanjurjo, A. (2006). Structural analysis of kasugamycin inhibition of translation. Nature Structural & Molecular Biology, 13(10), 879–886. https://doi.org/10.1038/nsmb1150

Seo, S. W., Kim, D., Szubin, R., & Palsson, B. O. (2015). Genome-wide Reconstruction of OxyR and SoxRS Transcriptional Regulatory Networks under Oxidative Stress in *Escherichia coli* K-12 MG1655. Cell Reports, 12(8), 1289–1299. https://doi.org/10.1016/j.celrep.2015.07.043

Shell, S. S., Wang, J., Lapierre, P., Mir, M., Chase, M. R., Pyle, M. M., Gawande, R., Ahmad, R., Sarracino, D. A., Ioerger, T. R., Fortune, S. M., Derbyshire, K. M., Wade, J. T., & Gray, T. A. (2015). Leaderless Transcripts and Small Proteins Are Common Features of the Mycobacterial Translational Landscape. PLOS Genetics, 11(11), e1005641. https://doi.org/10.1371/journal.pgen.1005641

Shenton, D., Smirnova, J. B., Selley, J. N., Carroll, K., Hubbard, S. J., Pavitt, G. D., Ashe, M. P., & Grant, C. M. (2006). Global Translational Responses to Oxidative Stress Impact upon Multiple Levels of Protein Synthesis. Journal of Biological Chemistry, 281(39), 29011–29021. https://doi.org/10.1074/jbc.M601545200

Slauch, J. M. (2011). How does the oxidative burst of macrophages kill bacteria? Still an open question: How do phagocytic ROS kill bacteria? Molecular Microbiology, 80(3), 580–583. https://doi.org/10.1111/j.1365-2958.2011.07612.x

Steiner, R. E., Kyle, A. M., & Ibba, M. (2019). Oxidation of phenylalanyl-tRNA synthetase positively regulates translational quality control. Proceedings of the National Academy of Sciences, 116(20), 10058–10063. https://doi.org/10.1073/pnas.1901634116

Tartaglia, L. A., Storz, G., & Ames, B. N. (1989). Identification and molecular analysis of oxyR-regulated promoters important for the bacterial adaptation to oxidative stress. Journal of Molecular Biology, 210(4), 709–719. https://doi.org/10.1016/0022-2836(89)90104-6

Temmel, H., Müller, C., Sauert, M., Vesper, O., Reiss, A., Popow, J., Martinez, J., & Moll, I. (2016). The RNA ligase RtcB reverses MazF-induced ribosome heterogeneity in *Escherichia coli*. Nucleic Acids Research, 45(8), 4708–4721. https://doi.org/10.1093/nar/gkw1018

Udagawa, T., Shimizu, Y., & Ueda, T. (2004). Evidence for the Translation Initiation of Leaderless mRNAs by the Intact 70 S Ribosome without Its Dissociation into Subunits in Eubacteria. Journal of Biological Chemistry, 279(10), 8539–8546. https://doi.org/10.1074/jbc.M308784200

Vesper, O., Amitai, S., Belitsky, M., Byrgazov, K., Kaberdina, A. C., Engelberg-Kulka, H., & Moll, I. (2011). Selective Translation of Leaderless mRNAs by Specialized Ribosomes Generated by MazF in Escherichia coli. Cell, 147(1), 147–157. https://doi.org/10.1016/j.cell.2011.07.047

Vila-Sanjurjo, A., Schuwirth, B.-S., Hau, C. W., & Cate, J. H. D. (2004). Structural basis for the control of translation initiation during stress. Nature Structural & Molecular Biology, 11(11), 1054–1059. https://doi.org/10.1038/nsmb850

Walz, A., Pirrotta, V., & Ineichen, K. (1976). λ represser regulates the switch between PR and Prm promoters. Nature, 262(5570), 665–669. https://doi.org/10.1038/262665a0

Willi, J., Küpfer, P., Evéquoz, D., Fernandez, G., Katz, A., Leumann, C., & Polacek, N. (2018). Oxidative stress damages rRNA inside the ribosome and differentially affects the catalytic center. Nucleic Acids Research, 46(4), 1945–1957. https://doi.org/10.1093/nar/gkx1308

Wu, J., Fan, Y., & Ling, J. (2014). Mechanism of oxidant-induced mistranslation by threonyl-tRNA synthetase. Nucleic Acids Research, 42(10), 6523–6531. https://doi.org/10.1093/nar/gku271

Zhang, Y., Zborníková, E., Rejman, D., & Gerdes, K. (2018). Novel (p)ppGpp Binding and Metabolizing Proteins of *Escherichia coli*. MBio, 9(2). https://doi.org/10.1128/mBio.02188-17

Zhong, J., Xiao, C., Gu, W., Du, G., Sun, X., He, Q.-Y., & Zhang, G. (2015). Transfer RNAs Mediate the Rapid Adaptation of *Escherichia coli* to Oxidative Stress. PLOS Genetics, 11(6), e1005302. https://doi.org/10.1371/journal.pgen.1005302

Zhu, M., & Dai, X. (2019). Maintenance of translational elongation rate underlies the survival of *Escherichia coli* during oxidative stress. Nucleic Acids Research, 47(14), 7592–7604. https://doi.org/10.1093/nar/gkz467

Zhu, Y. Y., Machleder, E. M., Chenchik, A., Li, R., & Siebert, P. D. (2001). Reverse transcriptase template switching: A SMART approach for full-length cDNA library construction. BioTechniques, 30(4), 892–897. https://doi.org/10.2144/01304pf02

