## Supplemental information for "Enhanced translation of leaderless mRNAs under oxidative stress in *Escherichia coli*"

### Supplementary information

*Supplementary Methods*

#### Strains and culture media

All experiments performed in this work used wild-type (WT) *E. coli* K-12 MG1655 strain. Strains were cultured in either lysogeny broth (LB) media (1% tryptone, 0.5% yeast extract, and 0.5% NaCl), or M9 media (47.7 mM $Na_2HPO_4$, 22.0 mM KH2PO4, 8.6 mM NaCl, 18.7 mM $NH_4Cl$, 2 mM $MgSO_4$, 0.1 mM $CaCl_2$, and 0.4% Glycerol). When indicated, branched amino acids (isoleucine, leucine, and valine 50 µg/ml each), 0.1% tryptone, 0.4% arabinose, 0.4%, ampicillin (100 µg/ml) or 250 µM paraquat were added to the culture media.

#### Analysis of polysomes

The detailed protocol for analysis of polysomes has been described by Qin and Kurt (2013) (Qin & Fredrick, 2013). In brief, 50 ml of M9 media supplemented with branched amino acids were inoculated with a saturated overnight culture in M9 media supplemented with tryptone and grown at 37°C in an orbital shaker. When indicated, the media also contained 250 µM paraquat. When bacteria reached mid-log phase (OD 600 ~0.4–0.6) the culture was poured on ice into a centrifuge bottle (250 mL) and the cells were spin down at 5000 RPM for 5 min. After discarding ice and supernatant, cells were resuspended in 50 µl lysis buffer (10 mM Tris-HCl, pH 8.0, 10 mM MgCl2, and 1 mg/ml lysozyme) and quickly frozen in liquid nitrogen. Frozen samples were thawed on ice and quickly frozen in liquid nitrogen three times. Samples were then thawed in the presence of 1/10 volume of 10% sodium deoxycholate, centrifuged at 5000 RPM for 5 min and loaded on a 10%-40% sucrose gradient. The gradients were centrifuged in a SW41 rotor at 35000 RPM for 3 hours and then analyzed using an in-line UV detector (ISCO/Brandel system).

#### Cloning and Mutation Protocols

Plasmid pBAD30SFIT (Rojas et al., 2018) contains a transcriptional fusion composed of superfold green fluorescent protein (sfGFP) and a modified mCherry, regulated by arabinose inducible BAD promoter (pBAD). The complete pBAD30SFIT was amplified using primers NotI-Fw and NotI-Rv to construct plasmid pSD by introducing a NotI restriction site upstream of the transcription start site of pBAD. Plasmid pSD was further modified to produce plasmid plmRNA by annealed oligo cloning in sites NotI-XhoI following previously established protocols (Normanly et al., 1986) and using oligonucleotides Oligo-lmGFP-Fw and Oligo-lmGFP-Rv. In the resulting plmRNA plasmid, the 5'UTR of *sfgfp* was removed.

To construct the plasmid pSD (*cI-gfp*), the first 100 codons of the leaderless *cI* transcription were cloned into NotI-XhoI restriction sites of plasmid pSD. After amplifying the gene fragment from λ phage genomic DNA using oligonucleotides cI-5'(PM)short-notI-Fw and cI-XhoI-Rv, the pcr product was digested using NotI and XhoI, followed by a ligation with the plasmid digested with the same restriction enzymes. This placed the start codon of *cI* next to the transcription start site.

A similar protocol was used to construct pSD (*katG(ACA)-gfp*) and pSD (*katG(AGA)-gfp*). After amplifying the 5' UTR and the first 100 *katG* codons from *E. coli* K-12 MG1655 genomic DNA using oligonucleotides katG(aca)-notI-Fw and katG-XhoI-Rv, the PCR product was digested and ligated into the NotI-XhoI restriction sites of plasmid pSD. To construct pSD (*katG(AGA)-gfp*), the protocol was repeated using oligonucleotides katG(aga)-notI-Fw and katG-XhoI-Rv, the latter of which contains the C→G point mutation.

*E. coli* K-12 MG1655 Δ*mazEF*::FRT was constructed by homologous recombination (Datsenko and Wanner, 2000) using plasmid pLZ01 (Blondel et al., 2013) as template for amplification of a Cam resistance cassette flanked by the FRT sites (FLP recombinase target sequence), as well as primers mazEF(H1+P2) and mazEF(H2+P1). Construction of *E. coli* K12 MG1655 Δ*relA*::FRT and Δ*lon*::FRT has been previously reported (Kelly et al., 2020). Deletion of *spoT* on the genome of the Δ*relA*::FRT strain was done following a similar protocol as for *MazEF* deletion, using primers spoT(H1+P1) and spoT(H2+P2). All deletions were transduced to the corresponding strain using phage P1vir.

All constructs and mutations were validated by Sanger sequencing.

All oligonucleotides mentioned in this protocol can be found at Supplementary Table S2.

**Translation Efficiency Analyses**

M9 media supplemented with branched amino acids and ampicillin were inoculated with bacteria from a saturated overnight culture in M9 media supplemented with tryptone and grown at 37°C in an orbital shaker. When the culture reached mid-log phase (OD 600 ~0.4–0.6), a 50 µl aliquot was diluted in a 96-well optical-bottom plate with 150 µl fresh M9 media supplemented with branched amino acids (50 µg/ml each) and arabinose (0.4% final concentration). When indicated, media additionally contained paraquat (250 µM final concentration). Plates were further shaken at 37°C. OD 600 and fluorescence intensity of green fluorescent protein (GFP; Ex. 480 ± 4.5 nm, Em. 515 ± 10 nm) and mCherry (Ex. 555 ± 4.5 nm, Em. 600 ± 10 nm) were measured in a microplate reader (Infinite M200 PRO, Tecan). As the ΔrelAΔspoT strain is unable to replicate in poor media, M9 was supplemented with 0.5% tryptone. When the culture reached mid-log phase, cells were washed twice with M9 media devoid of amino acids, before inoculating 96-well plates where it was supplemented with the branched amino acid mix or triptone as indicated in the text.

In all experimente, strains transformed with pBAD30 (parental plasmid of pBAD30SFIT) were used to subtract the fluorescence background.

**RNA extraction and RT-PCR**

M9 media supplemented with branched amino acids were inoculated with bacteria from a saturated overnight culture in M9 media supplemented with tryptone and grown at 37°C in an orbital shaker. 1 ml of bacterial cultures in M9 media supplemented with branched amino acids were pelleted for 1 min at 12,000 g. Pellet were resuspended in 50 µl lysis buffer (83 mM Tris HCl, pH 6.8, 18 mM EDTA pH 8.0, 1.7% SDS, and 1.6% 2-mercaptoethanol) and incubated for 3 min at 37°C. 1.0 ml of TRIzol (Ambion) was added, and total RNA was extracted following the manufacturer’s protocol.

RNA samples were treated with DNase I (Roche) and used to prepare cDNA using the RevertAid First Strand cDNA Synthesis Kit (Thermo), according to the manufacturer's protocols. Real-time PCR was performed using the SensiFAST™ SYBR® Hi-ROX kit (Bioline) and primers S7, X15, Y12 and pre16S for experiments shown in Figure 3 or primers sfgfp_5end_fw and sfgfp_5end_rv for region 1, mcherry_5end_fw and mcherry_5end_rv for region 2 and mcherry_3end_fw and mcherry_3end_rv for region 3 for experimentos shown in figure S1 (sequences at Supplementary Table S2).

**Determination of 5' end sequences of reporter transcripts**

The sequence of the 5' end of lmRNA and SD reporters was determined using the strategy published by Zhu *et al*. (Zhu *et al.*, 2001) . 1 mL aliquots of cultures in M9r media were pelleted for 1 min at 12,000 g. Pellets were resuspended in 50 µl lysis buffer (83 mM Tris HCl, pH 6.8, 18 mM EDTA pH 8, 1.7% SDS, and 1.6% 2-mercaptoethanol) and incubated for 3 min at 37°C. 1.5 ml of TRIzol was added, and total RNA was extracted following the manufacturer's protocol. RNA samples were treated with DNase I (Thermo), according to the manufacturer, to remove genomic and plasmid DNA. For first strand cDNA synthesis 3.5 µL of RNA (10-200 ng) were mixed with 1 µL of 25 mM primer S1sfGFP_3'XhoI_Rv, incubated at 65°C for 2 min and immediately cooled on ice. 4.5 µL of cold First-Strand reaction mix (2 µL of 5X First-Strand Buffer, 0.25 µl of 20 mM DTT, 1 µl of 10 mM dNTP mix, 0.25 µl of RiboLock RNase Inhibitor (40 U/µL) and 1 µl of SuperScript II Reverse Transcriptase (200U/µL)) were added. Mixture was incubated 3 min at 25°C, followed by 1 min at 42°C. 1 µL of 12 mM primer PP_TS was added and incubated 1 hour at 42°C. The cDNA was amplified by PCR using Taq Polymerase (Promega) using primers PP_A and S1sfGFP_3'XhoI_Rv, cloned on pGEMT-Easy, and sequenced using T7 and SP6 universal primers. Sequence of all primers can be found in Table S2.

*Supplementary Tables*

#### Table S1. 5' end of mRNA generated from the canonical and leaderless reporters.

| Reporter | Sequence*[1] |
|---|---|
| **pSD** | 5'-ATACCCGTTTTTTTGGGCTAGCGAATTCAGGAGGAATTTACC**<u>ATG</u>**AGCAAACTCGAGGGC... |
| **plmRNA** | 5'-ATA**<u>ATG</u>**AGCAAACTCGAGGGC... |

*[1] Sequence of the 5' end of pSD and plmRNA reporters as determined using the SMART method (Zhu et al., 2001) with RNA purified from bacteria cultured in the same conditions as the rest of the work. The GFP start codon is highlighted in bold and underlined.

#### Table S2: Oligonucleotides used in this work.

| Primer | Sequence* |
|---|---|
| NotI-fw | 5'-aaggaaaaaagcggccgctctctactgtttctccatacccgt |
| NotI-rv | 5'-aaggaaaaaagcggccgcaaaaagcgtcaggtaggatccgct |
| Oligo-lmGFP-Fw | 5'-ggccgctctctactgtttctccataatgagcaaac |
| Oligo-lmGFP-Rv | 5'-tcgagtttgctcattatggagaaacagtagagagc |
| cI-5'(PM)short-notI-Fw | 5'-ggaaaaaagcggccgctctctactgtttctccatatgagcacaaaaaagaaaccattaac |
| cI-XhoI-Rv | 5'-ccgctcgagacttctaagtgacggctgcatactaac |
| katG(aca)-notI-Fw | 5'-aaggaaaaaagcggccgctctctactgtttctccataacactgtagaggggagcacattg |
| katG(aga)-notI-Fw | 5'-aaggaaaaaagcggccgctctctactgtttctccataacactgtagaggggagcagattg |
| katG-XhoI-Rv | 5'-ccgctcgagaaacagaccggcgtaactgccccagtc |
| MazEF (H1+P2) | 5'-ctacatatgatagcggtttgaggaaagggttatgatccaccatatgaatatcctccttag |
| MazEF (H2+P1) | 5'-ctgtgaccagaatagaagtgagttagtaacactacccaatgtgcaggctggagctgcttc |
| SpoT (H1+P1) | 5'- cgggtcgcccttgtatctgtttgaaagcctgaatcaactggtgcaggctggagctgcttc |
| SpoT (H2+P2) | 5'- tcataaaacattaatttcggtttcgggtgactttaatcaccatatgaatatcctccttag |
| X15_rRNA | 5'-tacgacttcaccccagt |
| Y12_rRNA | 5'-taaggaggtgatccaaccgc |
| S7_rRNA | 5'-agaatgccacggtgaatacg |

| | |
|---|---|
| pre16S_rv | 5'-atcagacaatctgtgtgagcac |
| sfgfp_5end_fw | 5’-gtggagaagaacttttcactgga |
| sfgfp_5end_rv | 5’-agcatcaccttcaccctctc |
| mcherry_5end_fw | 5’-tgccgacattagaaatagcaca |
| mcherry_5end_rv | 5’-cgtgaccgttaacagaaccc |
| mcherry_3end_fw | 5’-ctggcgcgtacaatgtgaat |
| mcherry_3end_rv | 5’-atccatgccaccggtagaat |
| S1sfGFP_3'XhoI_Rv | 5'-tgcccattaacatcaccatc |
| PP_TS | 5'-caggacgctgttccgttctRaRuRgRgRg |
| PP_A | 5'-caggacgctgttccgttc-3' |

* Where in use, “R” indicates the utilization of ribonucleotides instead of deoxyribonucleotides in the next sequence position

**Supplementary Figures**

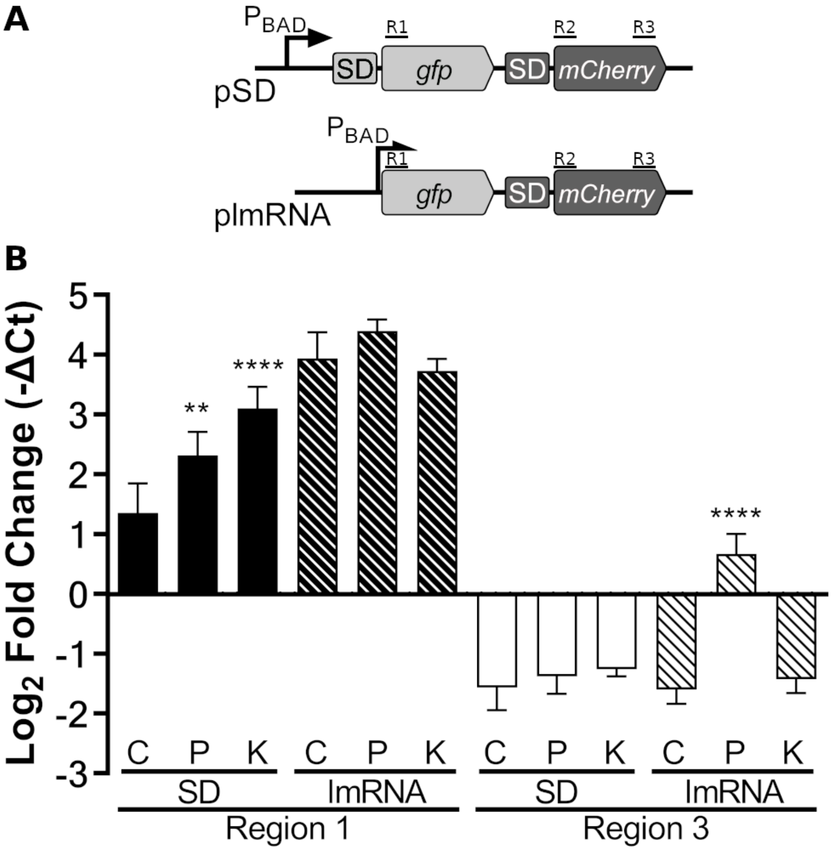


**Figure S1. Unbalanced RNA degradation does not explain increased GFP/mCherry fluorescence ratio.** A) Schematic representation of the PCR strategy used to analyze the integrity of the transcript ends. Using real-time PCR, levels of region 1 (R1) and region 3 (R3) were determined relative to region 2 (R2), for both pSD and plmRNA transcripts. B) The fold changes (-ΔCt) were determined under control condition (C) or stress induced with either 250 µM paraquat (P) or 750 µg/mL of kasugamycin (K). Statistical analyses: One-way ANOVA and Dunnett's multiple comparisons test. ****p<0,0001, **p<0,01.

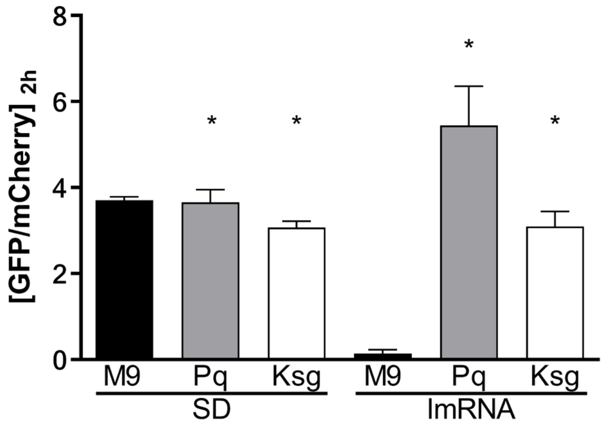


**Figure S2. Changes in lmRNA translation produced by kasugamycin can be followed by changes in GFP/mCherry fluorescence ratio.** GFP/mCherry fluorescence ratio for wild-type *E. coli* K-12 MG1655 carrying pSD and plmRNA reporters 2 h after transcription induction in control condition (M9) and under stress induced with either 250 µM Paraquat (Pq) or 750 µg/mL Kasugamycin (Ksg). Statistical analyses: Kruskal-Wallis test and Dunn's multiple comparisons test. *p<0,05.

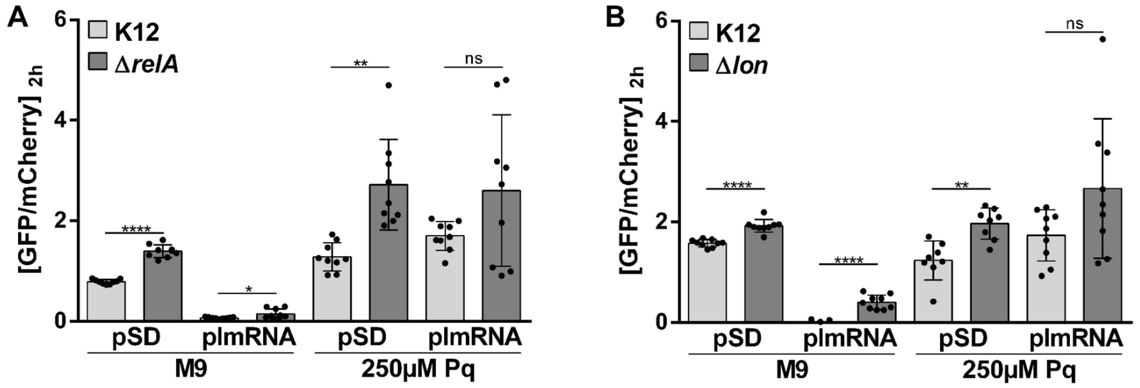


**Figure S3. Effects of *relA* or *lon* deletion on the translation of canonical and leaderless mRNA.** Comparison of the GFP/mCherry fluorescence ratio of pSD and plmRNA reporters at 2 h after transcription induction in control and stress conditions, in wild-type *E. coli* (K12) versus (A) *E. coli* Δ*relA*::FRT (Δ*relA*) and (B) *E. coli* Δlon::FRT (Δ*lon*). Statistical analyses: unpaired two-tailed t test with Welch's correction for A and B. ****p<0,0001, **p<0,01, *p<0,05.

***References of supplementary material***

Kelly, P., Kavoor, A., & Ibba, M. (2020). Fine-Tuning of Alanyl-tRNA Synthetase Quality Control Alleviates Global Dysregulation of the Proteome. *Genes*, *11*(10), 1222. https://doi.org/10.3390/genes11101222

Normanly, J., Masson, J. M., Kleina, L. G., Abelson, J., & Miller, J. H. (1986). Construction of two Escherichia coli amber suppressor genes: TRNAPheCUA and tRNACysCUA. *Proceedings of the National Academy of Sciences*, *83*(17), 6548–6552. https://doi.org/10.1073/pnas.83.17.6548

Qin, D., & Fredrick, K. (2013). Analysis of Polysomes from Bacteria. In *Methods in Enzymology* (Vol. 530, pp. 159–172). Elsevier. https://doi.org/10.1016/B978-0-12-420037-1.00008-7
